# A novel thin plate spline methodology to model tissue surfaces and quantify tumor cell invasion in organ-on-chip models

**DOI:** 10.1101/2023.11.20.567272

**Authors:** Elizabeth Elton, Carly Strelez, Nolan Ung, Rachel Perez, Kimya Ghaffarian, Naim Matasci, Shannon M. Mumenthaler

## Abstract

Organ-on-chip (OOC) models can be useful tools for cancer drug discovery. Advances in OOC technology have led to the development of more complex assays, yet analysis of these systems does not always account for these advancements, resulting in technical challenges. A challenging task in the analysis of these two-channel microfluidic models is to define the boundary between the channels so objects moving within and between channels can be quantified. We propose a novel imaging-based application of a thin plate spline method – a generalized cubic spline that can be used to model coordinate transformations – to model a tissue boundary and define compartments for quantification of invaded objects, representing the early steps in cancer metastasis. To evaluate its performance, we applied our analytical approach to an adapted OOC developed by Emulate, Inc., utilizing a two-channel system with endothelial cells in the bottom channel and colorectal cancer (CRC) patient-derived organoids (PDOs) in the top channel. Initial application and visualization of this method revealed boundary variations due to microscope stage tilt and ridge and valley-like contours in the endothelial tissue surface. The method was functionalized into a reproducible analytical process and web tool – the Chip Invasion and Contour Analysis (ChICA) – to model the endothelial surface and quantify invading tumor cells across multiple chips. To illustrate applicability of the analytical method, we applied the tool to CRC organoid-chips seeded with two different endothelial cell types and measured distinct variations in endothelial surfaces and tumor cell invasion dynamics. Since ChICA utilizes only positional data output from imaging software, the method is applicable to and agnostic of the imaging tool and image analysis system used. The novel thin plate spline method developed in ChICA can account for variation introduced in OOC manufacturing or during the experimental workflow, can quickly and accurately measure tumor cell invasion, and can be used to explore biological mechanisms in drug discovery.

## INTRODUCTION

Organ on a chip (OOC) models are *in vitro* systems meant to replicate morphological and functional properties of a specific organ or tissue (Leung 2022, Roth 2019). With respect to a diseased organ, such as cancer, OOC models allow for recapitulating aspects of the heterocellular tumor microenvironment (TME) and facilitate investigations of cell-cell interactions and cancer progression (Ahn 2017, Jouybar 2023, Strelez 2021). As a results, these models possess considerable value for mechanistic studies and drug discovery applications (Caballero 2017, Jouybar 2023). While OOC models present an exciting development in complex *in vitro* studies, analytical workflows to extract data from OOCs lag in development due to technical challenges, such as the 3D structures of the chips, multiple cell types cultured together, the small numbers of cells present in the models, and the inherent variability within these systems (Caballero 2017, Chaw 2007). Two significant drawbacks of OOC-based assays as currently performed involve single timepoint measurements taken in a disruptive fashion (i.e., endpoint assays) and frequently require bulk analyses of multiple heterogenous cell types. Improving these workflows to include spatial and cell type-specific information and to allow for dynamic measurements would support the use of OOCs for higher throughput experiments and aide drug discovery investigations. Imaging-based assays often provide this flexibility and resolution; however, challenges exist with OOC imaging and analysis due to the variable and complex nature of the devices. These difficulties are present across multiple OOC designs and make accurately identifying specific features, such as the tissue:tissue interface, challenging (Chen 2017). To account for these issues, we propose a novel application of a thin plate spline (TPS) methodology to represent tissue boundaries and quantify cancer cell dynamics in an analytical pipeline.

A TPS is a generalized cubic spline that can be used to model coordinate transformation and visualize model surfaces of varying smoothness (Gao 2010, Donato 2002, Wood 2003). TPS’s are regularly used to model surfaces in geosciences for climate, topographical, and oceanographic mapping (Hancock 2006, Trossman 2011). In the medical field, TPSs have been applied to ultrasound images of brain deformation, integrated into surgical tools for organ modelling with stereo imaging data, and applied to MRI data to search for deformations in heart muscle (Frisken 2019, Yang 2014, Amini 1998). As this method has been expanded to model many different types of surfaces or boundaries represented by highly variable and complex data, we hypothesized that a TPS method could alleviate some of the challenges related to OOC imaging and image analysis. In this study, we developed an analytical workflow, referred to as the topographical method, applied to image analysis outputs from a previously developed colorectal cancer (CRC) OOC model system. This model was combined with confocal microscopy to monitor CRC patient-derived organoid (PDO) cancer cell invasion from a top epithelial channel into a bottom endothelial channel, mimicking intravasation, an early step of cancer metastasis. Our method accounts for technical artifacts such as tilt variations in the microscope stage while also capturing fine details in the contours of the endothelial tissue, leading to a more accurate measurement of tumor cell behavior. This analytical workflow was developed into a web tool, the Chip Invasion and Contour Analysis (ChICA), to extract spatial information within OOC models and enable higher throughput analysis of invasion, making these chips more viable models for drug discovery and cancer research applications.

## METHODS

### CRC Organoid-on-Chip Imaging-based Invasion Assay

We utilized a previously described stretchable CRC organoid-on-a-chip comprised of a top epithelial compartment and a bottom endothelial compartment separated by a porous membrane (manufactured by Emulate, Inc., see Strelez 2021, 2023). Briefly, green fluorescent protein (GFP)-expressing CRC PDOs were seeded in the top channel and human umbilical vein endothelial cells (HUVECs) or human intestinal microvascular endothelial cells (HIMECs) labeled with red fluorescent protein (RFP) were seeded in the bottom channel. After six days, invasion from the epithelial compartment into the endothelial compartment was visualized by fluorescence confocal microscopy using the Operetta CLS High Content Analysis Platform (Revvity). Specifically, the co-culture region where the top and bottom channels overlap was imaged at 5 *μ*m intervals from the bottom of the endothelial compartment to the top of the epithelial compartment (50 slices) for a total z-height of 250 *μ*m. GFP+ tumor cells and RFP+ endothelial cells were segmented and identified based on florescence intensity using 3-D image analysis in the Revvity Harmony software. Additional processing in (shown in Supplemental Table 1) included removal of non-cell objects and selecting the endothelial cell population comprising the top layer of the endothelial compartment. Morphological and positional information for each object was exported for further analysis and downstream identification of invaded GFP+ tumor cells.

### Surface Modelling and Statistical Analysis

The positional output from Harmony image processing was used in a surface modelling and statistical analysis workflow to label GFP+ objects as ‘invading’ or ‘non-invading’. The data analysis workflow was developed using R Statistical Software (v4.2.2; R Core Team 2023) using the fastTps function from the fields package (v15.2; Nychka 2016). Using this function, we modelled the endothelial cell boundary layer as a thin plate spline. This technique, designed for use in topographical analysis, approximates the z coordinate of the endothelial cells as a function of the x and y coordinates of those cells. This package also allows for us to measure the standard error of the TPS model fit to the actual data. We calculated the confidence interval using the standard error to account for the thickness of the tissue, ensuring that actively invading epithelial cells that lie within the contours of the endothelial layer are accurately assigned invading status. Further quantification of the number of cells that are in the process of invading or have invaded is done by comparing the actual z-height coordinates of the epithelial cells to the predicted z-height based on the TPS-generated surface. If the actual z-height is below that of the surface, including the confidence interval, the cell is considered ‘invading’ and counted. Summary statistics of the ‘invading’ cells are generated using tidyverse packages (v2.0.0; Wickham 2019).

### Web Tool Development

The topographical workflow was developed into a web tool that was built using R Shiny. (Chang, 2023). The tool includes options for users to upload data, run the analysis, and visualize the surface and invading cells from each chip model. The included surface plotting function also utilizes tools from the fields R package to create an approximate surface of the endothelial landscape (v15.2; Nychka 2016). The 3D plotting tool reduces the coordinates from the spline to a 100 by 100 matrix for efficiency in plotting. The code used to build the web tool and generate figures can be found on GitHub at: https://github.com/eitm-org/chip_invasion

## RESULTS

### Higher-throughput imaging workflow for CRC organoid-on-chips

To establish an OOC imaging-based invasion workflow we implemented the following steps: (1) we imaged the overlapping epithelial and endothelial compartments of multiple CRC OOCs, (2) analyzed the positional and morphological data of tumor cells and endothelial cells, and (3) developed a standard R Shiny tool for tumor cell invasion quantification and visualization (Fig.1).

**Figure 1.**
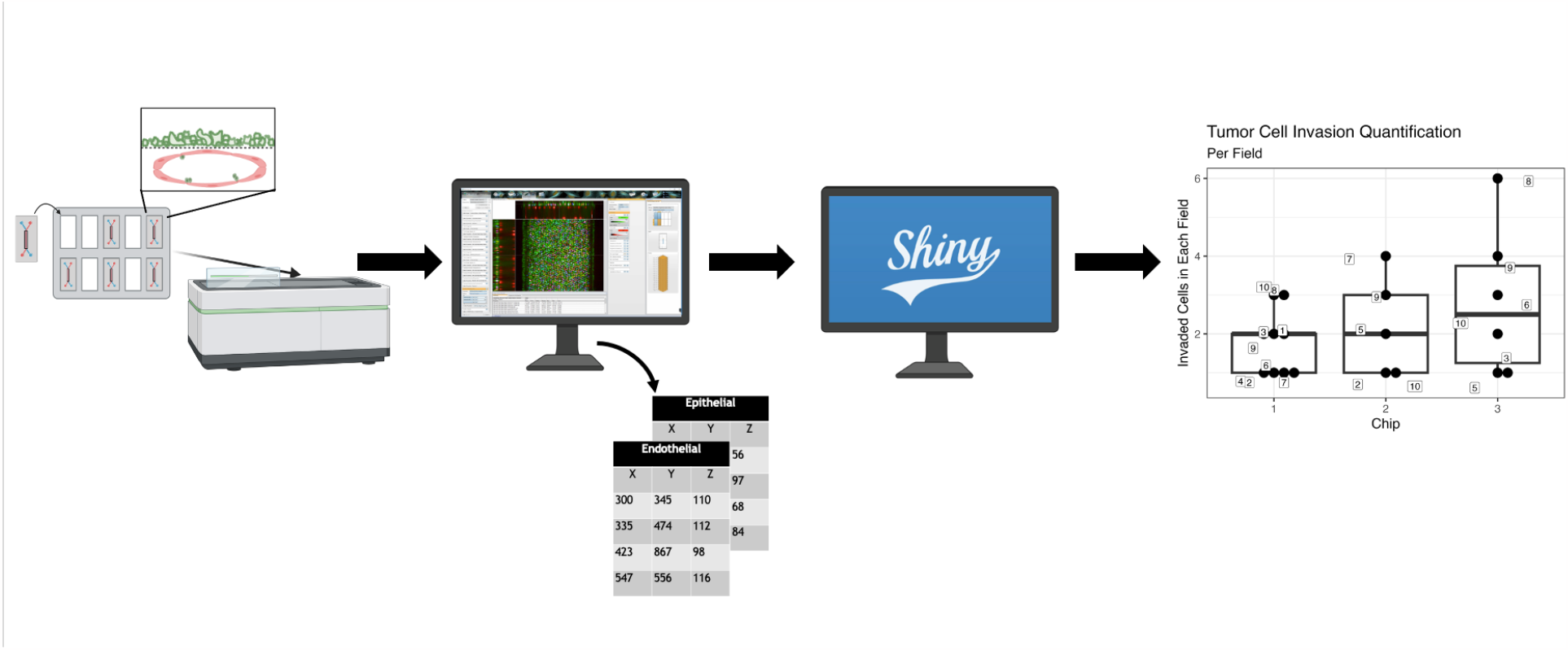
Workflow schematic. Multiple OOCs are seeded with GFP+ colorectal cancer patient-derived organoids in the top channel and RPF+ endothelial cells in the bottom channel and imaged on a high content imaging platform. Image analysis is performed to extract positional and morphological data for each cell type. Data analysis is performed using novel methodology and the Chip invasion and Contour Analysis web tool (ChICA) to quantify tumor cell invasion.

The workflow was built to analyze CRC-on-chip models comprised of CRC PDOs seeded into an extracellular matrix (ECM)-coated “top” channel and endothelial cells seeding into an ECM-coated “bottom” channel. OOCs were imaged at various time points (day 0 and day 6) and cyclic stretching was initiated to mimic the peristaltic forces of the intestine. OOCs were imaged along the length of the co-culture region from the bottom of the endothelial channel to the top of the epithelial channel (50 slices at a 5 *μ*m step size for a total Z-height of 250 *μ*m using the Operetta CLS High Content Analysis platform and pre-processed using Revvity Harmony image analysis software. This process included identifying GFP+ tumor cell objects and RFP+ endothelial cell objects and measuring morphological and spatial information, including x, y, and z position, intensity, and dimensions of the GFP+ and RFP+ objects. Detailed information about the Harmony analyses can be found in Supplemental Table 1. These images revealed a persistent tilt in the OOCs due to the alignment of chips in the stage adapter (Supplemental Fig. 1), slight variations in OOC manufacturing, and the presence of actively invading tumor cell objects, which could not be accounted for in image preprocessing. We therefore utilized the x, y, and z position of the GFP+ PDOs and RFP+ endothelial cells from Harmony image analysis to measure and define the boundaries between channels. The goal of this analysis was to accurately denote tumor cells as either ‘invading’ or ‘non-invading’ based on their presence in the top or bottom channel. To solve issues presented by manual quantification, we created a standardized and reproducible workflow in R.

### Thin plate spline for robust and reproducible surface modelling and tumor cell quantification

A thin plate spline was chosen to model the endothelial boundary based on its ability to account for tilt in any direction and smaller scale inconsistencies in the endothelial layer, as well as the ability to easily compare the relative position of other objects outside of the endothelial layer to the surface position (Boyd 1999). Initial application and visualization of this method confirmed the microscope stage tilt seen in the images and revealed that the endothelial tissue surface was not a flat plane but showed ridge and valley-like contours. 2D visualization of these endothelial surfaces also showed that while the presence of contours was consistent across chips, the shape, pattern, and frequency of the contours were heterogeneous (Fig. 2A). Next, the x and y coordinates of the GFP+ objects were fed into the spline to identify the invasion phenotype of the tumor objects. If the z-height was below the z-height predicted by the spline plus standard deviation of the spline, then the object was assigned ‘invading’ status. This method was compared to a simpler quantification method using a flat line method of mean endothelial Z-height plus standard deviation, which was found by manual assessment to overcount tumor cell objects in some fields of the chip and undercount in others due to the stage tilt. It also raised the concern that this method was not capturing actively invading objects, as image preprocessing indicated that these objects tended to sit within the contours of the endothelial layers. Compared to the global threshold, the topographical method captures actively invading cells that reside within the contours of the endothelial tissue regardless of tilt or other surface inconsistencies (Fig. 2B).

**Figure 2.**
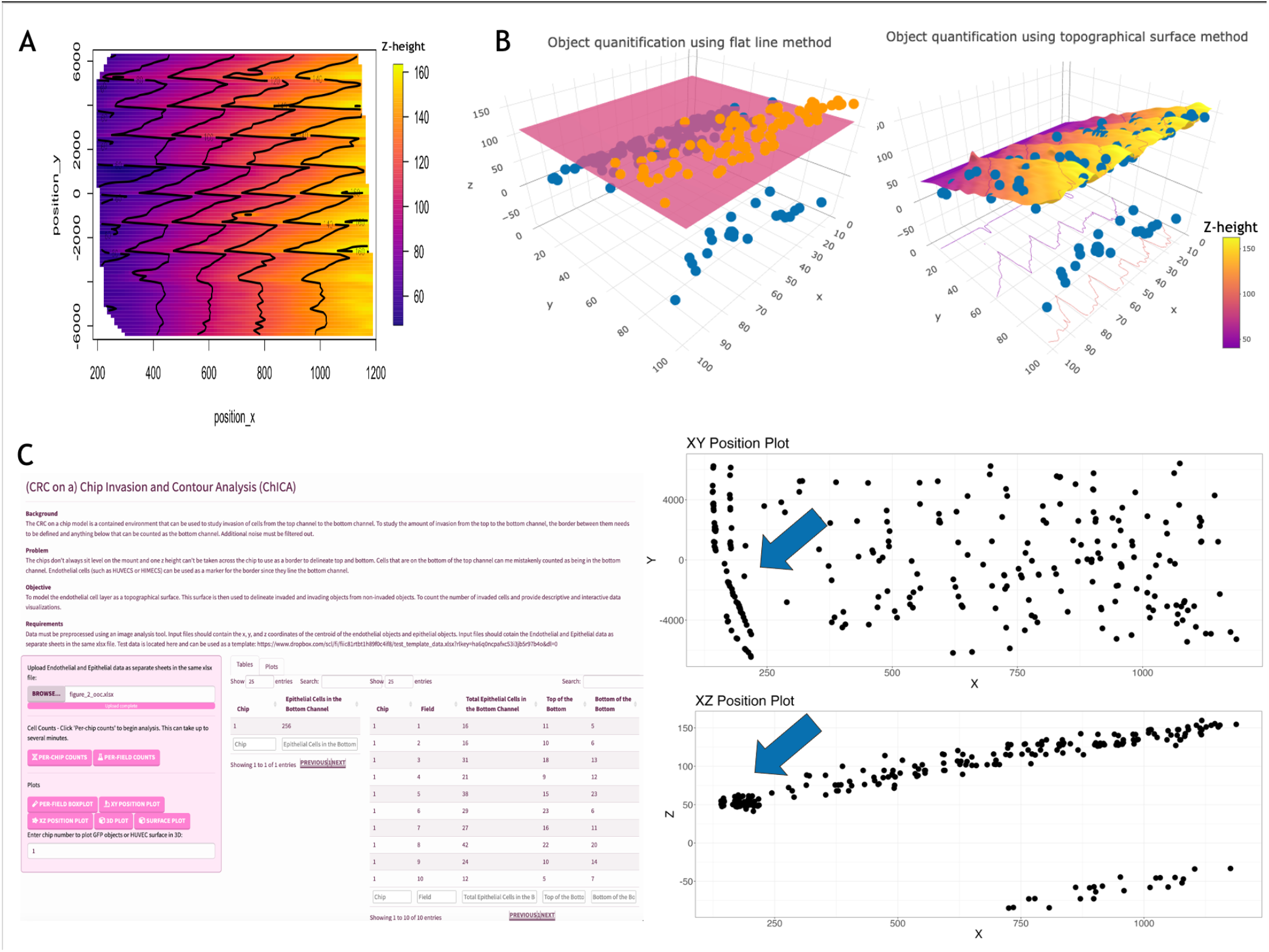
Endothelial surface modelling using topographical method enhances data extraction from high-throughput OOC experiments. (A) The topographical surface method is applied to an OOC seeded with HUVECs and imaged on day 0. A surface is generated by fitting a thin plate spline to positional endothelial data and is visualized in 2-D, with x and y position shown on the x and y axes and z-height shown using a gradual color scale. The surface demonstrates wave-like contours and a gradual decline in z-height from high x positions to low x positions. (B) The topographical surface method from A is compared to the use of a flat surface to quantify tumor invasion. In the flat line method (left), objects that are below the mean z-height plus standard error of the endothelial centroids are labelled as invading. The topographical method (right) labels objects as invading when they are below the endothelial surface plus standard error. In either method, objects labelled as invading are denoted in blue. Objects not labelled as invading are denoted in orange. (C) The topographical surface workflow was built into a publicly available web tool requiring no computational background to use. The user interface of the tool (left) provides options to upload data and visualize results in a table or graph form. The tool includes options to plot invaded tumor cell objects by their x and y position (right upper) and x and z position (right lower) for quality control. These plots captured invaded objects on the edge of the OOC model missed in image analysis quality control, indicated with arrows.

Once it was demonstrated that the topographical method could capture actively invading cells, the method was functionalized into a reproducible analytical process to read in the imaging results from multiple chips, model the endothelial surface, and quantify invading tumor cells. This workflow was implemented into a web tool – the Chip Invasion and Contour Analysis (ChICA) – to ensure reproducibility, allow for higher throughput analysis of multiple (6-12) chips at once, and expand accessibility of the workflow. Using ChICA, the x-y and x-z positions of the invaded objects were visualized for quality control. For example, in a OOC with slight manufacturing irregularities where the top and bottom channel did not completely overlap, the visualizations showed invaded tumor cell objects on the edge of the OOC where there were no endothelial cells present (Fig. 2C). Users could further refine the image analysis pipeline to exclude these fields to accurately assess tumor cell invasion in the presence of endothelial cells. After quality control is assessed, experimentalists can use the included template to upload the positional output of their invading epithelial objects and boundary endothelial objects. Positional data can come from any image analysis workflow, and any data preprocessing or cleaning work should be done during this step. Once uploaded, users can generate a surface model of their tissue, quantify the number of invaded objects, and visualize both the endothelial surface and invaded objects in 2-D and 3-D. The ChICA tool can be accessed at http://chica.eitm.org:8080/.

### ChICA allows for dynamic measurements of tumor organoid invasion on-chip

To illustrate functionality of the analytical method, we applied the tool to CRC organoid-chips seeded with different endothelial cell types and measured tumor cell invasion dynamics. Initially, HUVECs, an endothelial cell type derived from umbilical cord, were used to represent the endothelial tissue in the CRC model due to their prominent use in biological studies (Ewald, 2021). To better recapitulate the colon physiology, we also optimized the use of colon-specific HIMECs. Application of ChICA reveals qualitative differences in the contours present in the endothelial surface on day 6. HUVECs demonstrated more regular contours that showed a consistent wave shape with fewer endothelial objects sitting far above or below the central endothelial layer. On the other hand, HIMECs showed greater variation between chips, with some chips demonstrating flatter surfaces, dramatic wave patterns, or dips or bowing across the surface (Fig. 3A). As shown in Figure 3B, the ChICA tool created the surface boundary despite variations in the surface created by either endothelial cell type. The tool also revealed that, in addition to qualitative differences in the endothelial surface layer, the endothelial cell types led to different amounts of invaded tumor cells in the bottom channel on day 6, with more invasion observed in chips seeded with HUVECs compared to HIMECs (Fig. 3B). Furthermore, we quantified tumor cell invasion over the course of the six-day experiment, by imaging the same chips on day 0 and day 6. Invasion increased over time, irrespective of the endothelial cell type (Fig. 3C). The biological differences between HUVECs and HIMECs and the resulting impact on organoid invasion behavior is enabled by the ChICA tool.

**Figure 3.**
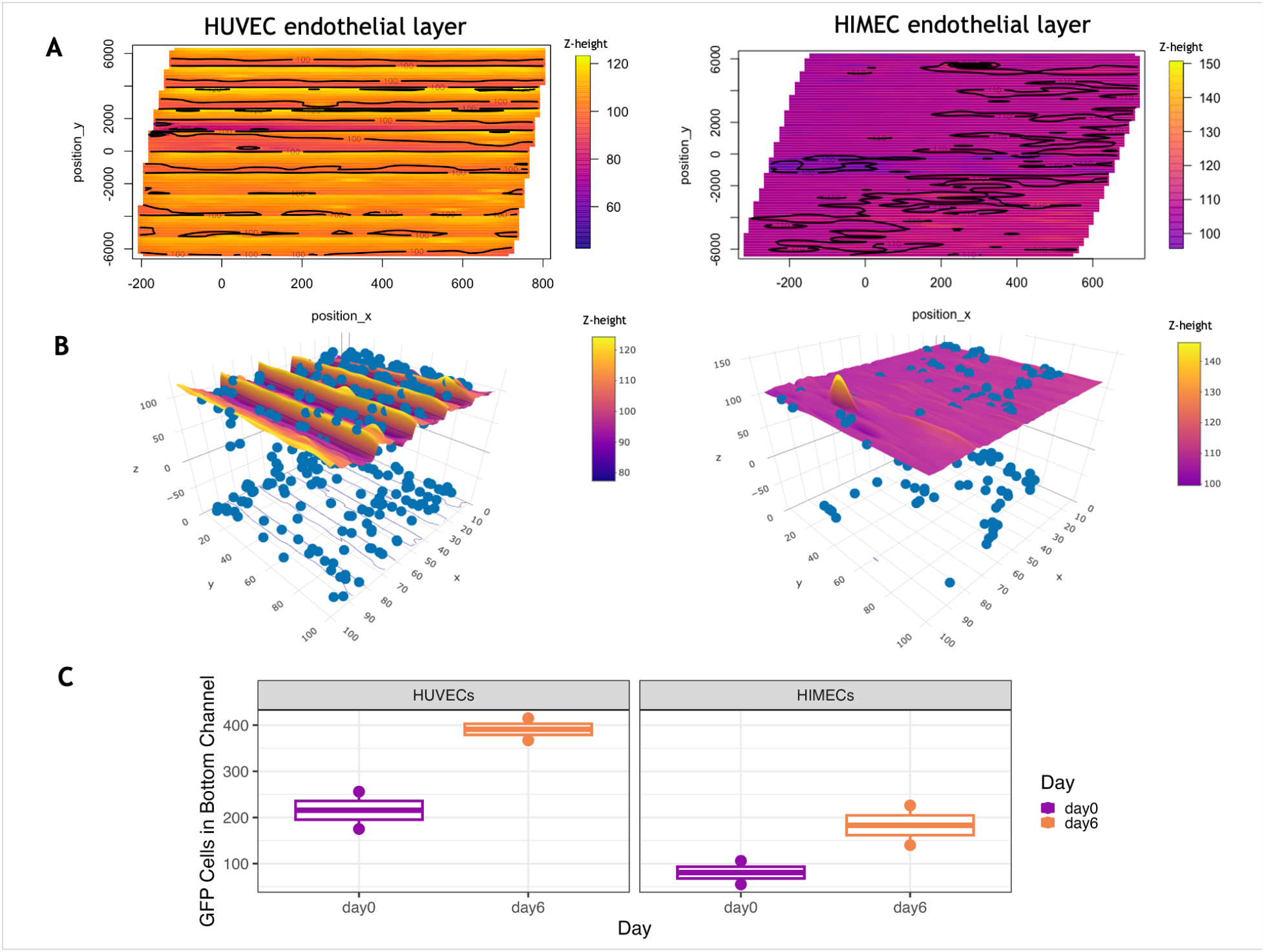
The topographical method labels and quantifies invading cells when applied to endothelial layers. (A) The topographical method is applied to OOCs with different endothelial cells and visualized in 2-D, with x and y position shown on the x and y axes and z-height shown using a gradual color scale. The surfaces formed from different endothelial cells demonstrate different contour patterns. The HUVEC endothelial surface (left) shows regular, wave-like contours of a similar height across the surface. The HIMEC endothelial surface (right) demonstrates irregular contours with occasional endothelial objects sitting above or below the center surface. (B) 3-D reconstructions of endothelial surfaces formed from HUVECs (left) or HIMECs (right) showing the boundary used to label tumor cell objects as invading or non-invading. The tumor cells are labeled blue. (C) Numbers of invaded GFP+ tumor cells as measured using the ChICA tool. CRC OOCs with either HUVECs (left) or HIMECs (right) in the endothelial compartment were imaged on day 0 and day 6 of the experiment and invaded cells were identified based on the relative position to the endothelial surface.

## DISCUSSION

We developed a web tool (ChICA) implementing a standardized analytical workflow based on a mathematical surface modelling technique to quantify tumor cell invasion in a CRC OOC model. This approach accounts for technical artifacts and reduces manual effort thus allowing for higher throughput and higher quality studies. While strides have been taken in the field of imaging to improve both the microscopes and their associated image analysis software, these tools often reduce analysis to a 2-D representation, introducing an opportunity to miss spatial features of the system (Peel 2021). Some open-source image analysis tools offer 3-D analysis options, but these tools often require computational expertise and depend on multiple manual filtering steps, limiting the ability to automate analysis and perform high throughput investigations (Kriston-Vizi, 2017). The ChICA web tool can be used to process multiple chips at once with a standard workflow, thus removing manual bias and increasing the speed at which researchers can extract meaningful information from imaging data. Furthermore, this tool allows users to investigate the 3D spatial dynamics of cells within the TME, including investigations of actively invading tumor cells – a critical population that might be missed by other approaches.

This approach provides accurate spatial information that can be used to elucidate the location of invading cells within the endothelial compartment (i.e., the top or bottom layer of the endothelial compartment), the presence of outlier objects, and capture phenotypic differences in the endothelial landscapes which might be associated with distinct cell types (e.g. HUVECs vs HIMECs) with biological or pharmacological relevance.

The methodology of representing the endothelial barrier as a contoured surface, rather than a globally flat boundary, is system agnostic and broadly applicable. Usage of the topographical workflow can be expanded to other cell types or diseases where cells move between compartments, such as monitoring immune cell trafficking, intravasation or extravasation in different diseased organs like cancers or irritable bowel disease. Other intestine or stomach-on-a-chip models, including those with villi or other boundary tissues with high levels of spatial variation, could benefit from high-fidelity 3-D reconstruction and corresponding data analysis workflow. Moreover, the ChICA tool can be used to quantify cell migration and viability readouts when multiple, distinct cell type are used and labeled.

Given the ChICA web tool requires only the coordinate information of the invading and boundary objects, we expect the approach to be broadly compatible with data from any OOC model, microscope, or image analysis software. Since the tool is downstream of any imaging and image preprocessing, the accuracy of the quantification is dependent on the quality of the image analysis and subsequent data cleaning. Users should ensure the validity of the classification and filtering features, which may depend on factors such as OOC design, cell size and fluorescent labeling, and the type of microscope, to get the most accurate surface model and cell counts. In future works, data cleaning features could be migrated from the image analysis software and into an R workflow to increase transparency and reproducibility.

The main advantage of using OOCs over simpler models is the preservation of the variation and heterogeneity of cell types and spatial organization present *in vivo*. Analytical workflows should account for this variation rather than reducing and masking the dimensionality of the system. ChICA exemplifies a data analysis workflow that not only accounts for the dimensionality of the model but uses it to its advantage. By acknowledging the spatial dynamics of the OOC model, drug discovery investigations can more effectively recapitulate how a biological system will respond to perturbations, enabling these investigations to be more informative.

## ACKNOWLEDGEMENTS

This research was supported by the NCI Tissue Engineering Consortium R01 CA241137 grant (SMM). We would like to thank Kian Kani for review of the manuscript, Abby Coleman for helpful discussions about mathematical representations, and Shiva Patre and Xingyao Chen for establishing a continuous integration and deployment platform for web tools.

