## Supplemental Figures for "A novel thin plate spline methodology to model tissue surfaces and quantify tumor cell invasion in organ-on-chip models"

Figure 1. Tilt in the microscope stage means that a global threshold does not accurately quantify invaded tumor cells

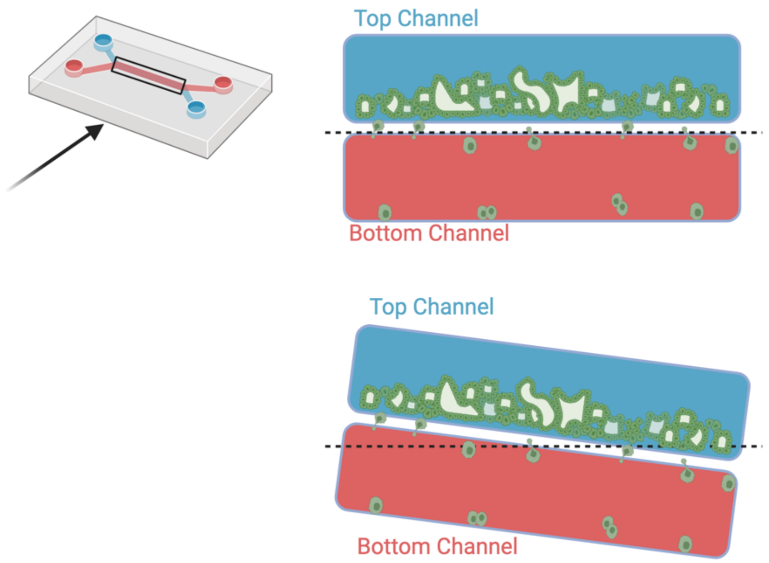


Table 1. Harmony Analysis Sequence

| Filtering Step | Parameters |
|---|---|
| Brightfield "Dirty/All" Binary | By Formula<br>Formula: A>100<br>Channel A: Brightfield<br>Negative Values: Set to Zero<br>Undefined Values: Set to Zero |
| Bright field - Filter Image - Texture PLS | Texture PLS<br>ROI: None<br>Filter: Plane Dark<br>Scale XY: 20 µm<br>Scale Z: 100 µm<br>PSF Aspect Ratio: 0 |
| Calculate image of the Brightfield Plane Dark in the previous image takes the B values greater than the B-mean multiplied by .5. | By Formula<br>Formula: B>B.mean*5<br>Channel B: Brightfield Plane Dark<br>Negative Values: Set to Zero<br>Undefined Values: Set to Zero |
| Calculate image was performed to subtract the two binary images, A=Brightfield "Dirty/All" Binary", B= Brightfield Plane Dark (PD) Binary | By Formula<br>Formula: A-B<br>Channel A: Brightfield "Dirty/All" Binary |

| | |
|---|---|
| | Channel B: Brightfield PD Binary (Outer Edge)<br>Negative Values Set to Zero<br>Undefined Values: Set to Zero |
| Find Image Region is performed on the Combined Binary previous image to find all regions using the absolute threshold with the lowest intensity >= 1 | Absolute Threshold<br>Lowest Intensity: ≥ 1<br>Highest Intensity: ≤ INF<br>Filling: Fill Plane-Wise<br>Volume: > 80000000 µm³ |
| Calculate Morphological Properties | Standard Volume |
| Filter Image - Searching for 546 Spot Bright (endothelial cells) location | Texture PLS<br>ROI: Chip Image Region<br>ROI Region: Chip Image Region<br>Filter: Spot Bright<br>Scale XY: 4 µm<br>Scale Z: 5 µm<br>PSF Aspect Ratio: 2 |
| Find Image Region (2) | Local Threshold<br>Threshold: 0.3<br>Region Scale: 25µm<br>Volume: >100µm³ |
| Calculate Position Properties | Standard Centroid Position in Image |
| Calculate Position Properties (2) | Z-Position of Centroid<br>Brightest Pixel: Alexa 546<br>Brightest Plane: Alexa 546 |
| Select Population | Filter by Property<br>546 Spot Bright Image Region Centroid Z [µm] :> 0 |
| Filter Image (3) | Sliding Parabola<br>Curvature: 100 |
| Filter Image (4) | Texture PLS<br>ROI: Chip Image Region<br>ROI Region: Chip Image Region<br>Filter: Spot Bright<br>Scale XY: 5 µm<br>Scale Z: 5 µm<br>PSF Aspect Ratio : 2 |
| Find Image Region (3) | Local Threshold<br>Threshold: 0.05<br>Region Scale: 15µm<br>Volume: > 500µm³ |
| Calculate Position Properties (3) | Standard Centroid Position in Image |
| Calculate Position Properties (4) | Z-Position of Centroid<br>Lowest/Middle/Highest Plane<br>Brightest Pixel: Alexa 488<br>Brightest Plane: Alexa 488 |

| | |
|---|---|
| Calculate Intensity Properties | Standard Mean<br>Standard Deviation<br>Coefficient of Variance<br>Quantile Fraction: 50 % |
| Calculate Morphology Properties (2) | Standard Volume<br>Surface Area<br>Equivalent Ellipsoid Axes<br>Object Box Size<br>Sphericity<br>Inner Sphere Radius<br>Object Height |
| Select Population (2) | Filter by Property<br>Intensity 488 Spot Bright Image Region<br>Alexa 488CV [%]: > 10<br>Intensity 488 Spot Bright Image Region<br>Alexa 488 StdDev: > 200<br>Boolean Operations: F1and F2 |
| **Object Results**<br>Population: Chip Image Region: ALL<br>Population: 546 Spot Bright Image Region: None<br>Population: 546 Upper Spot Bright: ALL<br>Population: 488 Spot Bright Image Region: None<br>Population: 488 Spot Bright Image Region Selected: ALL | |

6
